# Causal relationships between substance use and insomnia

**DOI:** 10.1101/2020.04.06.027003

**Authors:** Joëlle A. Pasman, Dirk J.A. Smit, Lilian Kingma, Jacqueline M. Vink, Jorien L. Treur, Karin J.H. Verweij

## Abstract

**Background:** Poor sleep quality and insomnia have been associated with the use of tobacco, alcohol, and cannabis, but it is unclear if there is a causal link. In this Mendelian Randomization (MR) study we examine if insomnia causes substance use and/or if substance use causes insomnia.

**Methods:** MR uses summary effect estimates from a genome-wide association study (GWAS) to create a genetic instrumental variable for a proposed ‘exposure’ variable and then identifies that same genetic instrument in an ‘outcome’ GWAS. With data of GWAS of insomnia, smoking (initiation, heaviness, cessation), alcohol use (drinks per week, dependence), and cannabis initiation, bi-directional causal effects were tested. Multiple sensitivity analyses were applied to assess the robustness of the findings.

**Results:** There was strong evidence for positive causal effects of insomnia on all substance use phenotypes (smoking traits, alcohol dependence, cannabis initiation), except alcohol per week. The effects on alcohol dependence and cannabis initiation were attenuated after filtering out pleiotropic SNPs. In the other direction, there was strong evidence that smoking initiation increased chances of insomnia (smoking heaviness and cessation could not be tested as exposures). We found no evidence that alcohol use per week, alcohol dependence, or cannabis initiation causally affect insomnia.

**Conclusions:** There were unidirectional effects of insomnia on alcohol dependence and cannabis initiation, and bidirectional effects between insomnia and smoking measures. Bidirectional effects between smoking and insomnia might give rise to a vicious circle. Future research should investigate if interventions aimed at insomnia are beneficial for substance use treatment.

## 1. Introduction

Insomnia (trouble falling and/or staying asleep) is associated with substance use, such as alcohol, cannabis, and nicotine use. Worldwide, individuals drink on average a glass of alcohol per day. A fifth of US and European adults smoke (WHO, 2016a), and a quarter to half of them have tried cannabis (EMCDDA, 2011). Both insomnia (Bin et al., 2012) and substance use (WHO 2016b, 2017, 2018) have serious consequences for work performance, risk of accidents, health, and well-being. Insight into the etiological processes underlying these associations might provide clues for prevention and intervention.

Use of alcohol, nicotine, and/or cannabis has been associated with increased prevalence of insomnia (Angarita et al., 2016; Sabanayagam and Shankar, 2011). These comorbidities may reflect a (partly) shared genetic etiology or causal relationships. For smoking, previous studies showed a genetic correlation with insomnia (Gibson et al., 2018; Jansen et al., 2019). As for causal relationships, experimental studies have investigated the acute effects of substance use on insomnia. Alcohol use shortened sleep onset latency, but led to sleep disruptions in the second half of sleep (Ebrahim et al., 2013). Cannabis intake likewise resulted in reduced sleep onset latency, but the effects of cannabis on sleep quality were less clear (Babson et al., 2017). Although smokers often cite its relaxing effects, nicotine intake was found to actually disturb sleep (Irish et al., 2015). Reversed causation -from insomnia to substance use-may also play a role. For example, adolescents with low sleep quality have shown a stronger inclination for later substance use (Hasler et al., 2016), although strong causal inferences cannot be made based on longitudinal designs.

Mendelian Randomization (MR) can be used for causal inference in complex relationships (Lawlor et al., 2008). A previous MR study found that insomnia increased smoking heaviness and decreased chances of cessation and found no effects in the other direction (Gibson et al., 2018). We extend this work by using genetic data from the largest genome-wide association studies (GWAS) to date to examine genetic and causal associations between insomnia and substance use, thereby also including alcohol and cannabis use.

## 2. Materials and methods

We first estimated genetic correlations between insomnia and substance use using LDscore regression (Bulik-Sullivan et al., 2015). Secondly, we used MR to test for causal effects of insomnia on substance use and vice versa. We used the Two-Sample MR R-package for the main analyses (Hemani et al., 2018) with GWAS summary statistics from non-overlapping samples (Table 1). The rationale behind MR is that genetic variants are randomly distributed in the population and therefore not affected by confounders. This makes genetic variants suited as instrumental variables to test causal effects of an ‘exposure’ on an ‘outcome’. Assumptions underlying MR are that the genetic instruments predict the exposure robustly (1) and are not independently associated with confounders (2) or the outcome (3). The latter two assumptions could be violated in case of horizontal pleiotropy (where one genetic variant directly affects multiple traits).

**Table 1.** Sources of the genome-wide association summary statistics used for the two-sample MR, the number of SNPs in the IVW exposure instrument (being the independent lead SNPs as reported in the source GWAS that were also present in the outcome SNP set, #exposure SNPs), the variance explained in the respective phenotype by these instrument SNPs (Instrument R^2^), and the genetic correlation of each substance use trait with insomnia (r_g_) with its associated *p*-value. For the computation of r_g_ we used the full GWAS summary statistics except for insomnia, where 23andMe participants were excluded.

| Phenotype | Source | Sample | #exposure SNPs<sup>a</sup> | Instrument $R^2$ | $r_g$, SE ($p$) |
| --- | --- | --- | --- | --- | --- |
| Insomnia | Jansen et al. (2019) | excl. 23andMe N=386,533 | 248 | 0.89% | NA |
| Smoking initiation | Liu et al. (2019) | excl. UKB* N=848,460 | 360 | 1.16% | .23, .02 (2.09E-23) |
| Smoking heaviness | Liu et al. (2019) | excl. UKB N=143,210 | NA<sup>a</sup> | NA<sup>a</sup> | .27, .03 (5.42E-17) |
| Smoking cessation | Liu et al. (2019) | excl. UKB N=216,590 | NA<sup>a</sup> | NA<sup>a</sup> | .28, .04 (5.56E-12) |
| Alcohol per week | Liu et al. (2019) | excl. UKB N=630,154 | 91 | 0.59% | .03, .02 (.029) |
| Alcohol dependence | Walters et al. (2018) | N=46,568 | 8 | 0.36% | .29, .07 (1.42E-5) |
| Cannabis initiation | Pasman et al. (2018) | excl. UKB N=57,980 | 32<sup>b</sup> | 1.33% | .04, .03 (.205) |

\*UKB=UK Biobank
<sup>a</sup> The effect of smoking heaviness and cessation on insomnia could not be tested because the insomnia GWAS could not be stratified on smoking status
<sup>b</sup> $p < 1e05$

Insomnia cases were people that reported they usually had trouble falling asleep at night or often woke up in the middle of the night; controls ‘never/rarely’ or ‘sometimes’ experienced this. Smoking initiation was defined as ever having smoked regularly (yes/no), smoking heaviness as cigarettes smoked per day, smoking cessation as having quit smoking (yes/no), and alcohol per week as the number of standard drinks consumed per week. Alcohol dependence (yes/no) was based on clinician’s diagnosis or on a semi-structured interview based on DSM-IV criteria. Cannabis initiation was defined as ever having used cannabis (yes/no). For the genetic instruments we selected independent SNPs that were genome-wide significantly associated with the exposure variable as reported in the source GWAS (p<5E-8; Table 1). However, for cannabis initiation there were only 2 genome-wide significant variants after excluding the UK Biobank sample, so for this phenotype we included SNPs that reached a ‘suggestive’ threshold of p<1E-5. Smoking heaviness and cessation could not be used as exposures, as the insomnia summary statistics could not be stratified on smoking status. Instrument strength was estimated by summing the variance explained (R^2^) by each independent instrument SNP in the exposure (Table 1).

The main analysis was an inverse-variance weighted (IVW) meta-analysis of the SNP-outcome association divided by the SNP-exposure association for each SNP. Sensitivity analyses were used to assess the robustness of the IVW findings against violation of the MR assumptions. Weighted median and weighted mode regression correct for effect size outliers that could represent pleiotropic effects (Hartwig et al., 2017). MR-Egger regression provides an intercept that indicates the presence of pleiotropy, and adjusts the regression coefficient for such effects (Bowden et al., 2015). MR-Egger relies on the NO Measurement Error (NOME) assumption, violation of which can be tested with the I^2^-statistic. When I^2^ was between 0.6 and 0.9, simulation extrapolation (SIMEX) was used to correct the MR-Egger for NOME violation; if I^2^ was below 0.6 MR-Egger was not reported (Bowden et al., 2016). We also applied Generalised summary-data-based MR (GSMR), which gains statistical power by taking small levels of linkage disequilibrium between the included SNPs into account, and deletes effect size outliers. Using Steiger filtering, MR analyses were repeated excluding SNPs that explained more variance in the outcome than in the exposure, and again retaining only SNPs that explained significantly (p<.05) more variance in the exposure (to rule out reverse causation; Hemani et al., 2017). Cochran’s Q-statistic was used to check for SNP effect heterogeneity (Bowden et al., 2018) and the F-statistic for weak instrument bias (Burgess et al., 2011). Finally, leave-one-out IVW analyses were used to give an indication of disproportional effects of single SNPs (Hemani et al., 2018). Rather than assessing the strength of the statistical evidence by p-values only, we also consider the effect sizes of the IVW and the sensitivity analyses to inform our interpretation.

## 3. Results

There were moderate genetic correlations between the full insomnia GWAS and all substance use GWAS except alcohol per week (small overlap) and cannabis initiation (no significant overlap; Table 1).

### Insomnia to substance use

The IVW analyses showed strong evidence for causal effects of insomnia on all substance use traits except alcohol use per week. For all analyses except insomnia-on-cannabis initiation there was evidence for SNP-effect heterogeneity, although leave-one-out analyses did not show the effects were driven by a single SNP (Figures S1-S6). The insomnia instrument had low explained variance, but did not suffer from weak instrument bias (F>10). I^2^, F, and Q statistics are presented in Table S1. The effect of insomnia on substance use retained similar effect sizes across the weighted mode, median, and GSMR analyses (although effect estimates became less precise; Table 2). MR-Egger results were not reported because the I^2^ statistic was below 0.6. For the smoking and alcohol use per week outcomes the proportion of SNPs that explained more variance in the outcome than in insomnia varied from 5.4 to 20.6% (Table S2). The Steiger-filtered IVW with those SNPs excluded showed only slightly attenuated effect sizes. However, when retaining only SNPs that significantly explained more variance in insomnia than in the outcome, strong evidence remained only for an effect on smoking initiation. For alcohol dependence (36.0%) and cannabis initiation (25.4%) large proportions of SNPs explained more variance in the outcome than in the exposure. Filtering those out led to substantial attenuation of the effects (Table 2).

**Table 2.**
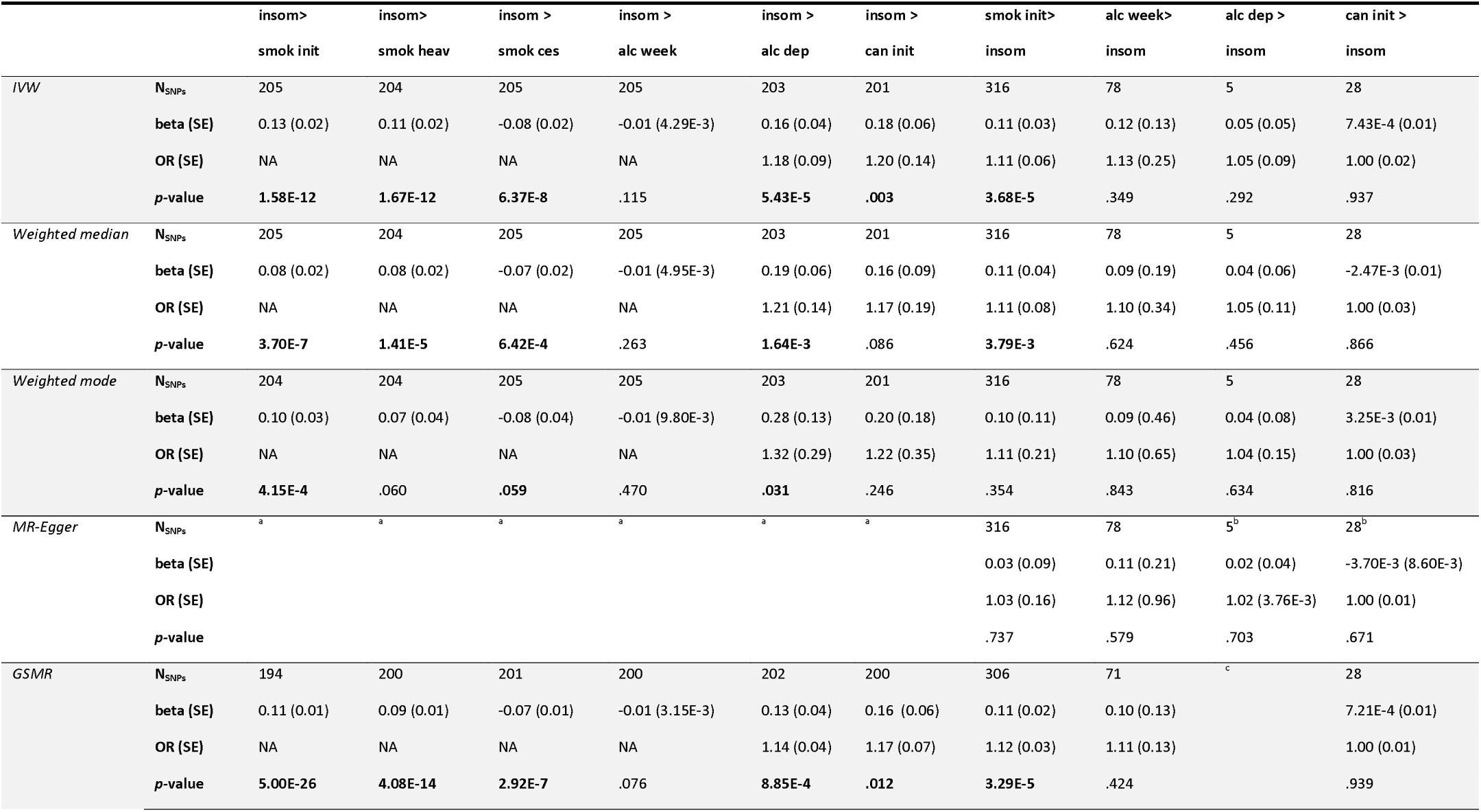
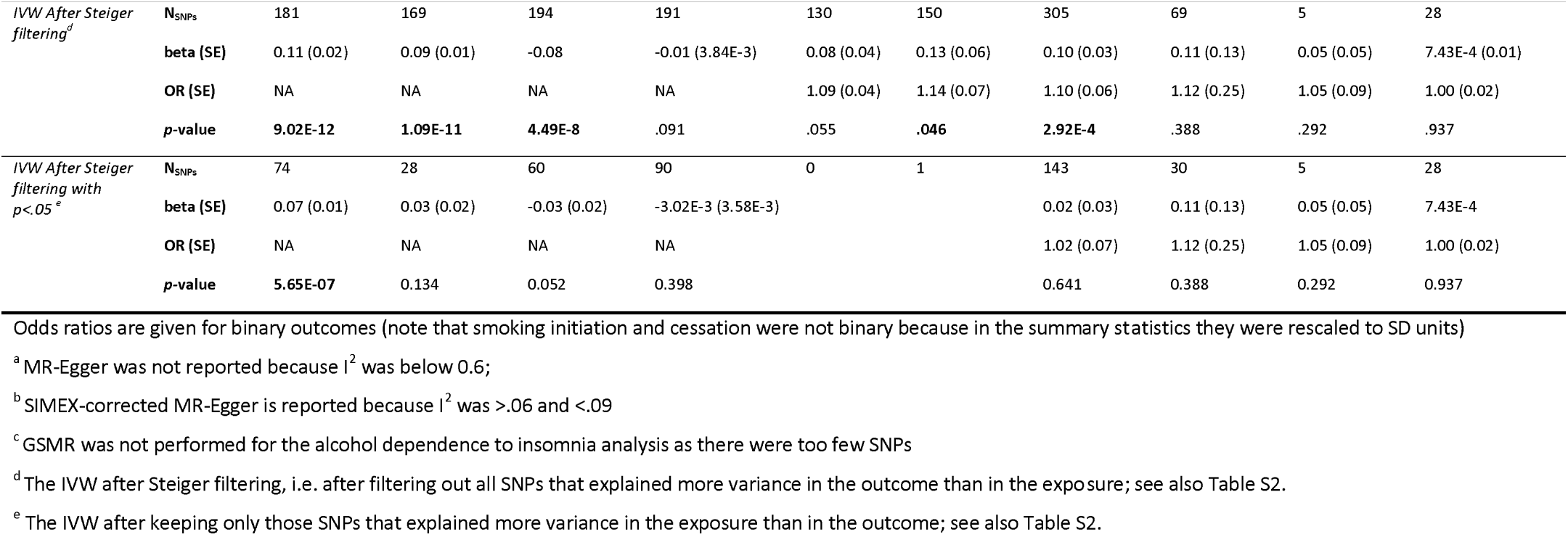
Results for the MR analyses with the IVW representing the main analysis and the remaining representing the results for the sensitivity analyses.

|  |  | insom><br>smok init | insom><br>smok heav | insom ><br>smok ces | insom ><br>alc week | insom ><br>alc dep | insom ><br>can init | smok init><br>insom | alc week><br>insom | alc dep ><br>insom | can init ><br>insom |
| --- | --- | --- | --- | --- | --- | --- | --- | --- | --- | --- | --- |
| *IVW* | **N<sub>SNPs</sub>** | 205 | 204 | 205 | 205 | 203 | 201 | 316 | 78 | 5 | 28 |
|  | **beta (SE)** | 0.13 (0.02) | 0.11 (0.02) | -0.08 (0.02) | -0.01 (4.29E-3) | 0.16 (0.04) | 0.18 (0.06) | 0.11 (0.03) | 0.12 (0.13) | 0.05 (0.05) | 7.43E-4 (0.01) |
|  | **OR (SE)** | NA | NA | NA | NA | 1.18 (0.09) | 1.20 (0.14) | 1.11 (0.06) | 1.13 (0.25) | 1.05 (0.09) | 1.00 (0.02) |
|  | **p-value** | 1.58E-12 | 1.67E-12 | 6.37E-8 | .115 | 5.43E-5 | .003 | 3.68E-5 | .349 | .292 | .937 |
| *Weighted median* | **N<sub>SNPs</sub>** | 205 | 204 | 205 | 205 | 203 | 201 | 316 | 78 | 5 | 28 |
|  | **beta (SE)** | 0.08 (0.02) | 0.08 (0.02) | -0.07 (0.02) | -0.01 (4.95E-3) | 0.19 (0.06) | 0.16 (0.09) | 0.11 (0.04) | 0.09 (0.19) | 0.04 (0.06) | -2.47E-3 (0.01) |
|  | **OR (SE)** | NA | NA | NA | NA | 1.21 (0.14) | 1.17 (0.19) | 1.11 (0.08) | 1.10 (0.34) | 1.05 (0.11) | 1.00 (0.03) |
|  | **p-value** | 3.70E-7 | 1.41E-5 | 6.42E-4 | .263 | 1.64E-3 | .086 | 3.79E-3 | .624 | .456 | .866 |
| *Weighted mode* | **N<sub>SNPs</sub>** | 204 | 204 | 205 | 205 | 203 | 201 | 316 | 78 | 5 | 28 |
|  | **beta (SE)** | 0.10 (0.03) | 0.07 (0.04) | -0.08 (0.04) | -0.01 (9.80E-3) | 0.28 (0.13) | 0.20 (0.18) | 0.10 (0.11) | 0.09 (0.46) | 0.04 (0.08) | 3.25E-3 (0.01) |
|  | **OR (SE)** | NA | NA | NA | NA | 1.32 (0.29) | 1.22 (0.35) | 1.11 (0.21) | 1.10 (0.65) | 1.04 (0.15) | 1.00 (0.03) |
|  | **p-value** | 4.15E-4 | .060 | .059 | .470 | .031 | .246 | .354 | .843 | .634 | .816 |
| *MR-Egger* | **N<sub>SNPs</sub>** | <sup>a</sup> | <sup>a</sup> | <sup>a</sup> | <sup>a</sup> | <sup>a</sup> | <sup>a</sup> | 316 | 78 | 5<sup>b</sup> | 28<sup>b</sup> |
|  | **beta (SE)** |  |  |  |  |  |  | 0.03 (0.09) | 0.11 (0.21) | 0.02 (0.04) | -3.70E-3 (8.60E-3) |
|  | **OR (SE)** |  |  |  |  |  |  | 1.03 (0.16) | 1.12 (0.96) | 1.02 (3.76E-3) | 1.00 (0.01) |
|  | **p-value** |  |  |  |  |  |  | .737 | .579 | .703 | .671 |
| *GSMR* | **N<sub>SNPs</sub>** | 194 | 200 | 201 | 200 | 202 | 200 | 306 | 71 | <sup>c</sup> | 28 |
|  | **beta (SE)** | 0.11 (0.01) | 0.09 (0.01) | -0.07 (0.01) | -0.01 (3.15E-3) | 0.13 (0.04) | 0.16 (0.06) | 0.11 (0.02) | 0.10 (0.13) |  | 7.21E-4 (0.01) |
|  | **OR (SE)** | NA | NA | NA | NA | 1.14 (0.04) | 1.17 (0.07) | 1.12 (0.03) | 1.11 (0.13) |  | 1.00 (0.01) |
|  | **p-value** | 5.00E-26 | 4.08E-14 | 2.92E-7 | .076 | 8.85E-4 | .012 | 3.29E-5 | .424 |  | .939 |
| *IVW After Steiger filtering<sup>d</sup>* | **N<sub>SNPs</sub>** | 181 | 169 | 194 | 191 | 130 | 150 | 305 | 69 | 5 | 28 |
|  | **beta (SE)** | 0.11 (0.02) | 0.09 (0.01) | -0.08 | -0.01 (3.84E-3) | 0.08 (0.04) | 0.13 (0.06) | 0.10 (0.03) | 0.11 (0.13) | 0.05 (0.05) | 7.43E-4 (0.01) |
|  | **OR (SE)** | NA | NA | NA | NA | 1.09 (0.04) | 1.14 (0.07) | 1.10 (0.06) | 1.12 (0.25) | 1.05 (0.09) | 1.00 (0.02) |
|  | **p-value** | 9.02E-12 | 1.09E-11 | 4.49E-8 | .091 | .055 | .046 | 2.92E-4 | .388 | .292 | .937 |
| *IVW After Steiger filtering with p<.05<sup>e</sup>* | **N<sub>SNPs</sub>** | 74 | 28 | 60 | 90 | 0 | 1 | 143 | 30 | 5 | 28 |
|  | **beta (SE)** | 0.07 (0.01) | 0.03 (0.02) | -0.03 (0.02) | -3.02E-3 (3.58E-3) |  |  | 0.02 (0.03) | 0.11 (0.13) | 0.05 (0.05) | 7.43E-4 |
|  | **OR (SE)** | NA | NA | NA | NA |  |  | 1.02 (0.07) | 1.12 (0.25) | 1.05 (0.09) | 1.00 (0.02) |
|  | **p-value** | 5.65E-07 | 0.134 | 0.052 | 0.398 |  |  | 0.641 | 0.388 | 0.292 | 0.937 |

Odds ratios are given for binary outcomes (note that smoking initiation and cessation were not binary because in the summary statistics they were rescaled to SD units)
<sup>a</sup> MR-Egger was not reported because $I^2$ was below 0.6;
<sup>b</sup> SIMEX-corrected MR-Egger is reported because $I^2$ was >.06 and <.09
<sup>c</sup> GSMR was not performed for the alcohol dependence to insomnia analysis as there were too few SNPs
<sup>d</sup> The IVW after Steiger filtering, i.e. after filtering out all SNPs that explained more variance in the outcome than in the exposure; see also Table S2.
<sup>e</sup> The IVW after keeping only those SNPs that explained more variance in the exposure than in the outcome; see also Table S2.

### Substance use to insomnia

The IVW analyses showed a causal effect of smoking initiation on insomnia, and no effects of the other traits. In the weighted median, mode, and GSMR sensitivity analyses the effect size of smoking initiation was roughly equal, although statistical evidence was slightly weaker (and substantially weaker in the weighted mode). Smoking initiation-on-insomia was the only analysis with sufficiently high I^2^ to allow for MR-Egger intercept interpretation, showing no evidence for pleiotropy (p=.347), even though the MR-Egger estimate was substantially attenuated. Less than 4% of the instrument SNPs explained more variance in insomnia outcome than in smoking initiation (Table S2). Filtering those out hardly changed results, although retaining only SNPs that explained significantly more variance in the exposure did substantially attenuate the effects (Table 2). There was no evidence for heterogeneity or weak instrument bias (Table S1, Figures S7-S10).

## 4. Discussion

There were moderate genetic correlations between insomnia and smoking initiation, smoking heaviness, smoking cessation, and alcohol dependence, such that insomnia was genetically associated with higher levels of substance use. The genetic correlation with alcohol per week was small but significant, and there was no significant correlation with cannabis initiation.

Overall, we found more evidence for causal effects from insomnia to substance use than vice versa. MR results suggest that insomnia leads to heavier smoking, increased chances of smoking initiation, alcohol dependence, and cannabis initiation, and decreased chances of smoking cessation. The finding that insomnia caused heavier smoking and lowers chances of smoking cessation confirms results from Jansen et al. (2019) and Gibson et al. (2018) on smoking and extends it to other substance use traits. As a possible interpretation, a desire to smoke may be induced by sleep deprivation (Hamidovic and de Wit, 2009). The causal effects of insomnia on alcohol use may be interpreted in light of a self-medication framework, as alcohol has somnolent properties (Goodhines et al., 2019). For cannabis the same reasoning might apply, although this interpretation seems more likely for a measure of cannabis use frequency rather than lifetime use. While we found an effect of insomnia on alcohol dependence, we found no effect on alcohol use. This might be due to the measure of alcohol use in quantity per week, which does not distinguish drinking large quantities in one evening from drinking one glass with dinner daily; the first would impair sleep quality more than the latter. The genetic architecture of drinking frequency seems to differ from that of drinking quantity (Marees et al., 2019).

In the other direction, we only found an effect of smoking initiation on insomnia. A previous study testing this relationship did not find this effect, possibly due to lower power (Gibson et al., 2018). The effect of smoking on insomnia might be due to nicotine’s stimulant properties (Greenland et al., 1998), although we could not test the effect of smoking heaviness. The absence of an effect of alcohol use and dependence on insomnia is in contrast with previous literature that suggested a negative effect of alcohol on sleep quality using experimental designs (Ebrahim et al., 2013). Our results might be due to the low instrument strength for the alcohol phenotypes. Also, the genetic instruments capture lifetime vulnerability to alcohol use and dependence, which is not directly comparable to the immediate effects of alcohol tested in experiments.

This study had both strengths and limitations. Results were reasonably robust against MR assumption violation. However, the effects of insomnia on alcohol dependence and cannabis initiation were in part driven by pleiotropic SNPs, suggesting caution in interpreting these findings. Although the analysis of smoking initiation on insomnia did not show strong evidence for it, pleiotropy might also play a role in this association. For example, smoking initiation has been linked to ADHD liability in children that have not started smoking yet, indicating that it might represent something different than only the inclination to use substances (Treur et al., 2019). Another limitation might be the use of instruments that explained limited amounts of variance in their respective phenotype (0.36-1.33%). Sensitivity analyses correcting for this showed attenuation in effect sizes. Finally, for cannabis initiation we used a more inclusive p-value threshold, which might have increased chances of pleiotropy. However, filtering out instruments that explained more variance in the exposure than the outcome did not have strong effects on the results.

To summarize, we find genetic overlap between insomnia and substance use, evidence for causal effects from insomnia to most substance use traits, and a causal effect of smoking initiation on insomnia. Future research should focus on underlying mechanisms and potential implications for clinical practice. As has been found previously (Patterson et al., 2017), our results suggest that treatment for substance use and insomnia could be optimized when attention is devoted to both issues.

## Abbreviations

insom: insomnia
smok init: smoking initiation
smok heav: smoking heaviness
smok ces: smoking cessation
alc week: alcohol use per week
alc dep: alcohol dependence
can init: cannabis initiation
IVW: inverse variance weighted meta-analysis
OR: odds ratio
GSMR: generalised summary-data-based Mendelian randomization
N_SNPs_: number of SNPs that was retained in the analyses after filtering for high LD, palindromic, and ambiguous SNPs, with additional HEIDI filtering in the GSMR and filtering for pleiotropic SNPs in the Steiger analyses.

## Acknowledgements

JLT is supported by a Veni grant from the Netherlands Organization for Scientific Research (NWO; grant number 016.Veni.195.016). KJHV and JLT are supported by the Foundation Volksbond Rotterdam. We would like to thank the research participants and employees of 23andMe for making this work possible.

